# Temporal coordination of collective migration and lumen formation by antagonism between two nuclear receptors

**DOI:** 10.1101/2020.03.16.993279

**Authors:** Xianping Wang, Heng Wang, Lin Liu, Sheng Li, Gregory Emery, Jiong Chen

## Abstract

During development, cells often undergo multiple, distinct morphogenetic processes to form a tissue or organ, but how their temporal order and time interval are determined remain poorly understood. Here we show that the nuclear receptors E75 and DHR3 regulate the temporal order and time interval between the collective migration and lumen formation of a coherent group of about 8 cells called border cells during *Drosophila* oogenesis. In wild type egg chambers, border cells need to first collectively migrate to the anterior border of oocyte before undergoing lumen formation to form micropyle, the structure that is essential for sperm entry into the oocyte. We show that E75 is required for border cell migration and it antagonizes the activity of DHR3, which is necessary and sufficient for the subsequent lumen formation during micropyle formation. Furthermore, *E75*’s loss of function or *DHR3* overexpression each leads to precocious lumen formation before collective migration, an incorrect temporal order for the two morphogenetic processes. Interestingly, both E75 and DHR3’s levels are simultaneously elevated in response to signaling from the EcR, a steroid hormone receptor that initiates border cell migration. Subsequently, the decrease of E75 levels in response to decreased EcR signaling leads to the de-repression of DHR3’s activity and hence switch-on of lumen formation, contributing to the regulation of time interval between collective migration and micropyle formation.

## Introduction

During development, a group or population of cells often has to undergo multiple, distinct morphogenetic processes in a certain temporal order (e.g. A, then B…) to form a tissue or organ (Webb and Oates, 2016). If the correct temporal order is not followed (e.g. process B occurring before process A), that tissue or organ would not form correctly (Rougvie, 2001; Thummel, 2001). Besides the correct order, the time interval between two processes is another important aspect of the temporal control for the morphogenetic processes. Making the interval too long or too short would also be detrimental to the formation of the organ or tissue. Despite their importance in development, our current understandings on how the temporal order and time intervals are regulated and determined still remain very limited.

The somatic follicle cells of the *Drosophila* egg chamber have served as an excellent model system to study multiple morphogenetic processes (Horne-Badovinac and Bilder, 2005). Specifically, during stage 9 of oogenesis, a group of about 8 cells detaches from the anterior follicle epithelium and undergoes collective migration between the germ-line nurse cells in a posterior direction (Montell, 2003). By early stage 10a, this coherent cluster of cells would have migrated a distance of about 150 μm in 6 hours, reaching the border between oocyte and nurse cells, hence the name border cells. About four hours later, by stage 10b, the cluster of 8 border cells would have migrated dorsally a short distance along the border, eventually stopping at the dorsal-most position of the border. Three hours later, by stage 12 or 13, this border cell cluster undergoes a second morphogenetic process to eventually form the tip of micropyle, a tubular structure required for sperm entry into the mature oocyte (Montell et al., 1992). Therefore, the formation of micropyle tip by border cells requires two distinct morphogenetic processes in a certain temporal order, first the well-studied, stereotyped, collective migration process and then a largely uncharacterized morphogenetic process that transforms these border cells into the tip of the tubular structure. Furthermore, an interval of about 16 hours exists between the beginning of collective migration and the start of the micropyle formation (Horne-Badovinac and Bilder, 2005). However, whether and how the temporal order and the time interval between the two morphogenetic processes are regulated remain largely unknown.

Previous studies have shed light on the temporal regulation of border cell migration. The steroid hormone ecdysone, its receptor heterodimer Ecdysone Receptor (EcR) and Ultraspiracle (USP), and their co-activator Taiman (Tai) had all been shown to be required for the initiation of border cell migration (Bai et al., 2000; Jang et al., 2009). Ecdysone and the EcR signaling had long been known to play important roles in coordination of growth and developmental timing during embryogenesis, larval molting and metamorphosis in *Drosophila* (Jia et al., 2017; Kozlova and Thummel, 2003; Yamanaka et al., 2013). Active form of ecdysone is also made in the adult *Drosophila* ovaries to regulate progression of oogenesis (Buszczak et al., 1999; Carney and Bender, 2000). 20-hydroxyecdysone, the active form of ecdysone, is locally synthesized by the follicle epithelium in individual egg chambers and reaches its highest levels around stages 9 and 10 (Domanitskaya et al., 2014; Margaret B et al., 1989). Even small patches of wild type follicle cells in mosaic stage 9 egg chambers were shown to produce a sufficient level of active ecdysone that allows the border cells to begin migration (Domanitskaya et al., 2014). The sufficiency of ecdysone and EcR signaling on initiation of border cell migration was further demonstrated by Jang and coworkers, in which early expression of the activated form of the co-activator Tai can precociously initiate border cell migration (Jang et al., 2009). However, what cellular processes in the border cells are directly regulated by EcR signaling and whether EcR also temporally regulates micropyle formation are currently unknown.

In this study, we show that E75 and DHR3, two nuclear receptors/transcription factors downstream of EcR signaling, regulate both the temporal order and time interval between border cell migration and micropyle formation. During border cell migration, EcR signaling activates the expression of both E75 and DHR3, with E75 repressing DHR3’s function. Furthermore, de-repression of DHR3 function after completion of border cell migration switches on lumen formation, turning the cluster of border cells into the tip of micropyle. Such antagonistic relationship between E75 and DHR3 (while both under the control of EcR signaling) provides the regulatory mechanism of temporal order and time interval between two distinct morphogenetic processes essential for the formation of a functional micropyle.

## Results

### RNAi Screen identifies E75 acting downstream of EcR signaling

Ecdysone signaling was known to be critical for the temporal control of initiation of border cell migration (Bai et al., 2000; Jang et al., 2009), but the cellular processes directly regulated by EcR signaling are largely unknown. To identify these, we carried out a small-scale RNAi screen of candidate genes that were previously reported to be responsive to ecdysone in *Drosophila* larvae and pupae and in cell lines (Beckstead et al., 2005; Gauhar et al., 2009; Sap et al., 2015). We first screened through the well-established response genes of ecdysone signaling (Ashburner, 1976; Huet et al., 1995; Yamanaka et al., 2013), including *E74, E75, E93, Br-c* and *DHR3*. Two to three different RNAi lines for each gene were used to confirm that phenotypes were not due to off-target effects, and two RNAi lines for the *EcR* gene were used as positive controls. A border-cells specific Gal4 driver, *Slbo-Gal4*, was used to drive expression of various *RNAi* transgenes in border cells beginning at late stage 8 of oogenesis, before border cells initiate their migration at early stage 9. As expected, both *EcR RNAi* lines (9327 and v35078 lines) resulted in phenotypes of strong migration delay or block, consistent with the previous reported roles of EcR in initiating and promoting border cell migration (Figures 1A, 1B and S1A-S1C) (Hackney et al., 2007; Jang et al., 2009). In comparison, border cell clusters within the wild type control stage 10 egg chambers almost always reached the 100% migration position, with only 6% of clusters displaying moderate delay (stopping at 75% migration position) (Figures 1B and S1A-S1C). Interestingly, of all the five ecdysone response genes tested, only *E75* displayed strong migration defects (Figures 1B and S1C). In fact, all three RNAi lines (v44851, 26717, Thu1738) consistently resulted in severe migration block and delay phenotypes, as compared to the control (Figures S1A and S1C). We then screened an additional collection of 20 genes that were considered ecdysone response genes or putative target genes of EcR/USP in recent reports (Beckstead et al., 2005; Gauhar et al., 2009; Li and White, 2003). However, none of the genes, when knocked down, displayed strong migration defects (Figure S1C). Only mild to moderate migration phenotypes were observed in a few of the RNAi experiments.

**Figure 1.**
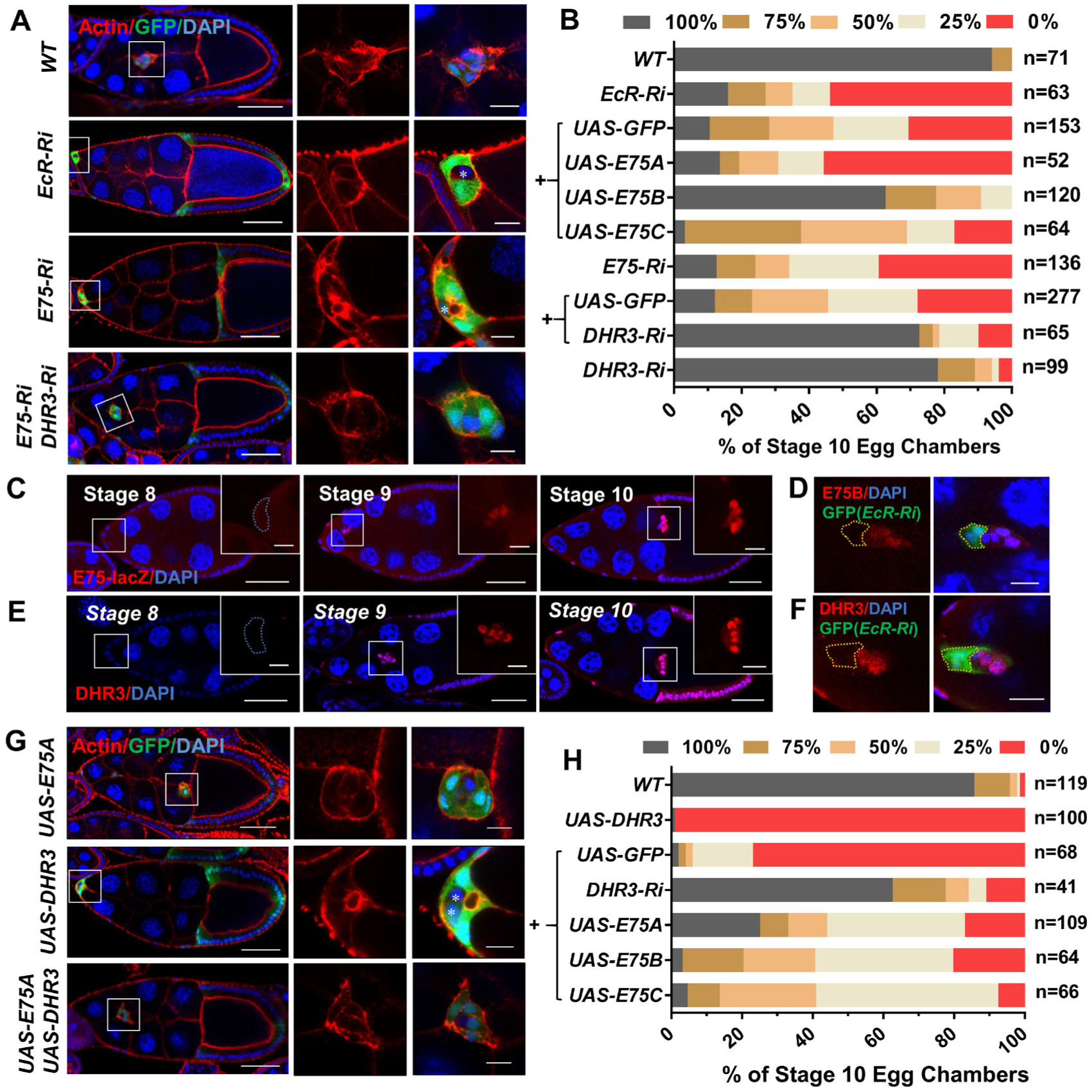
E75 antagonizes DHR3 during border cell migration. (A) Confocal images of egg chambers stained with phalloidin (red, for F-actin) and DAPI (blue, for nuclei) with indicated genotypes. The boxed regions are enlarged and shown to the right. Border cells expressing *EcR RNAi* displayed strong migration defects but exhibited similar morphology and F-actin distribution pattern to those of wild type (WT) border cells, whereas *E75 RNAi* border cells with migration defects displayed different morphology and F-actin distribution pattern from those of the *EcR RNAi* and WT border cells. Co-expression of *DHR3 RNAi* rescued *E75 RNAi*’s morphology and F-actin defects. *Ri* is the abbreviation for *RNAi* for this and all subsequent figures. Posterior is to the right and anterior is to the left for this and all subsequent figures. (B) Quantification of border cell migration with indicated genotypes. *EcR-Ri* denotes *EcR RNAi*, and”+”indicates that these genotypes include both *EcR RNAi* and one of the denoted genotypes (*UAS-GFP, UAS-E75A, UAS-E75B*, and *UAS-E75C*). Th”e +”below *E75 RNAi* indicates that these genotypes includes both *E75 RNAi* and one of the denoted genotypes (*UAS-GFP, DHR3 RNAi*). The stock used for *E75 RNAi* is v44851, which is used for all the other experiments unless noted otherwise. The x-axis denotes the percentage of stage 10 egg chambers examined for each genotype that exhibited various degrees of migration, as represented by the five color-coded bars (see Figures S1B and S1C for details). The 100% migration category (grey) indicates completion of migration, whereas 0% (red) indicates severe migration block. And the 25%, 50% and 75% categories indicate various degrees of migration delay. (C, E) Confocal images displaying β-galactosidase staining (C) and DHR3 staining (E) of stages 8, 9 and 10 egg chambers. Boxed region is enlarged to the right, showing a high-magnification view of the border cells. (D, F) Confocal images showing antibody staining of E75B (D) and DHR3 (F) of individual stage 10 border cell clusters with flip-out clones expressing *EcR RNAi* (*EcR-Ri*). The flip-out clones (labeled by GFP and encircled by yellow dotted lines) clearly displayed marked reduction of E75B and DHR3 respectively. (G) Border cells overexpressing *DHR3* exhibited severe defects in migration and morphology, which could be rescued by co-expression of *E75A*. Border cells with *E75A* overexpression alone displayed wild type phenotype. * (in A and G) indicates polar cells that are labeled by absence of GFP. (H) Quantification of rescue of border cell migration defects as resulted from *DHR3* overexpression. “+”indicates that these genotypes include both *UAS-DHR3* and one of the denoted genotypes (*UAS-GFP, DHR3 RNAi, UAS-E75A, UAS-E75B*, and *UAS-E75C*). Scale bars: 50 μm in (A, C, E, G), 10 μm for high-magnification views in (A, C-F, G). See also Figures S1 and S2.

We then proceeded to determine whether E75 acts downstream of EcR to initiate and promote border cell migration. Three distinct isoforms of E75 (A, B, C) were shown to be involved in different developmental and cellular processes and manifested stage- and tissue-specific responses (Li et al., 2016; Terashima and Bownes, 2006), and the sequences used in the three RNAi lines for *E75* are all within the common region and would have knocked down all three isoforms. Therefore, we overexpressed each isoform to test its individual rescue ability on border cell migration defects that were caused by *EcR RNAi*. We found that *E75B* overexpression markedly rescued *EcR RNAi*’s migration defects, whereas *E75C* displayed a much weaker rescue effect and *E75A* showing no significant rescue (Figure 1B). Moreover, we found that *E75*’s overall transcription levels (as represented by a previously used reporter *E75-lacZ* (Manning et al., 2017) within border cells at stages 9 and 10 were much higher than those at stage 8 (Figure 1C), consistent with ecdysone signaling being significantly increased beginning at stage 9. And mosaic border cell clusters containing a clone of *EcR RNAi* expressing cells demonstrate that E75B protein levels are drastically decreased when EcR function is reduced (Figure 1D), indicating that EcR activity is required for *E75B* expression during stage 9. Taken together, these results demonstrate that E75B is the major downstream player, among the previously known ecdysone response genes, to mediate EcR’s temporal control on border cell migration. Consistently, a recent study using microarray analysis also identified *E75* as a target gene that is responsive to ecdysone signaling in the migratory border cells (Manning et al., 2017).

### E75 antagonizes DHR3’s function during collective migration of border cells

During metamorphosis, ecdysone-activated EcR turns on the expression of E75B, which then binds to DHR3 and antagonizes its activity (White et al., 1997). E75B and DHR3 are both nuclear receptors/transcription factors and are both induced by ecdysone, and E75B’s inhibition of DHR3 function leads to suppression of DHR3’s transcriptional activation of its target genes essential for metamorphosis (Caceres et al., 2011; Reinking et al., 2005; White et al., 1997). To determine whether antagonistic interaction also exists between E75 and DHR3 during border cell migration, we co-expressed *DHR3 RNAi* and *E75 RNAi* in border cells. We found that DHR3 reduction strongly rescued *E75 RNAi*’s migration defects (Figures 1A and 1B), as well as the morphological defects of border cells (Figure 1A, also described in the section below). On the other hand, overexpression of *DHR3* resulted in similar phenotypes of migration and morphology to those of *E75 RNAi* (Figures 1G and 1H), with *DHR3* overexpression’s defects more severe than those of *E75 RNAi* (Figures 1B, 1H, S3A and S3B). Furthermore, *E75* overexpression can in turn suppress *DHR3* overexpression’s severe defects (Figures 1G and 1H), with all three of its isoforms (E75A, E75B, E75C) displaying similar suppressing abilities. This is consistent with previous reports that both E75A and E75B isoforms can heterodimerize with DHR3 to inhibit DHR3’s transcription activation ability (Sullivan and Thummel, 2003; White et al., 1997). Lastly, we showed that DHR3’s levels were also increased in border cells beginning at stage 9 (Figure 1E), similar to E75’s temporal expression pattern (Figure 1C), and its levels also depended on EcR’s activity (Figure 1F). Together, these data demonstrate an antagonistic relationship between E75 and DHR3 during border cell migration, with both their expressions activated by EcR during the migratory process.

Conversely, we found that expressing *E75 RNAi* (stock 26717) or *DHR3* in border cells significantly reduced the level of ecdysone response activity or EcR signaling (Figures S2A-S2C and S2F), which is represented by expression levels of *EcRE-lacZ*, a common reporter of EcR activity used in previous studies (Jang et al., 2009; Koelle et al., 1991). Moreover, expression of *DHR3 RNAi* could rescue *E75 RNAi*’s *EcRE-lacZ* expression levels (Figures S2D-S2F). These data indicate that E75 can exert a positive feedback on EcR signaling by antagonizing DHR3’s inhibition effect on EcR signaling. This conclusion is consistent with previous studies that showed DHR3 physically interacted with EcR and suppressed its activity (Lam et al., 1997; White et al., 1997). These results suggest that one of the means that E75 mediates EcR’s migration-promoting function is through E75’s positive feedback on EcR signaling.

### E75 antagonizes DHR3’s function in lumen formation during border cell migration

We noted that border cell clusters with *E75* knockdown or *DHR3* overexpression displayed different morphology and F-actin staining pattern from border cells with reduced EcR function (Figures 1A and 1G), as indicated by our *EcR RNAi* result and previous reports (Hackney et al., 2007; Jang et al., 2009). The delayed border cell clusters with *EcR RNAi* often displayed a coherent and front-polarized morphology with F-actin enriched in the front periphery of the cluster, similar to that of the wild type clusters (Figure 1A). On the contrary, *E75 RNAi* or *DHR3* overexpressing border cells lost the front-polarized cluster morphology that is characteristic of front-back polarity, and F-actin is instead enriched in the center of the cluster in a ring-like structure (Figures 1A and 1G), which is unique and never observed in any of the previously reported mutant phenotypes of border cells (to our knowledge). Closer examination revealed that this unique structure is not within individual border cell’s cytoplasm but is instead composed of portions of outer border cells’ inside membranes, which are joined together to form a continuous supra-cellular ring (Figures 2A-2E). Moreover, this supra-cellular structure is also enriched with molecules that are typically associated with apical membranes (aPKC, Crb, Baz/Par3, PIP2-GFP reporter) (Figures 2A, 2C and 2E) but not with lateral membranes (Dlg) (Figures 2A). A typical supra-cellular ring encloses a space that resembles a lumen with significant depth (about 5-10 μm, Movie S1) in the center of cluster, effectively displacing the two central polar cells to the side and underneath (Figures 2A, 1A and 1G; marked by *). The strong and specific enrichment of apical markers such as aPKC in the membranes enclosing the luminal space suggests that the border cell cluster has undergone a lumen formation process to become a tubular structure with the apical membrane facing the central lumen. Interestingly, the *E75 RNAi* border cells displayed a range of lumen-like phenotypes. Half of them (50.0%) showed a clear lumen phenotype that is similar to that of the *DHR3* overexpressing border cells, while majority of the rest (39.0%) exhibited little luminal space and discontinuous apical membrane patches as labeled by aPKC (Figure S3A and S3B), which resemble the previously reported structure of pre-apical patches (PAP) that are present during the intermediate stages of *de novo* lumen formation in several model systems (Bryant et al., 2010; Ferrari et al., 2008; Yang et al., 2013). These moderate phenotypes may reflect incomplete lumen formation or the intermediate stages of lumen formation in the border cells, while the large lumen structure from almost all of the *DHR3* overexpressing border cells and half of the *E75 RNAi* border cells may indicate complete lumen formation.

**Figure 2.**
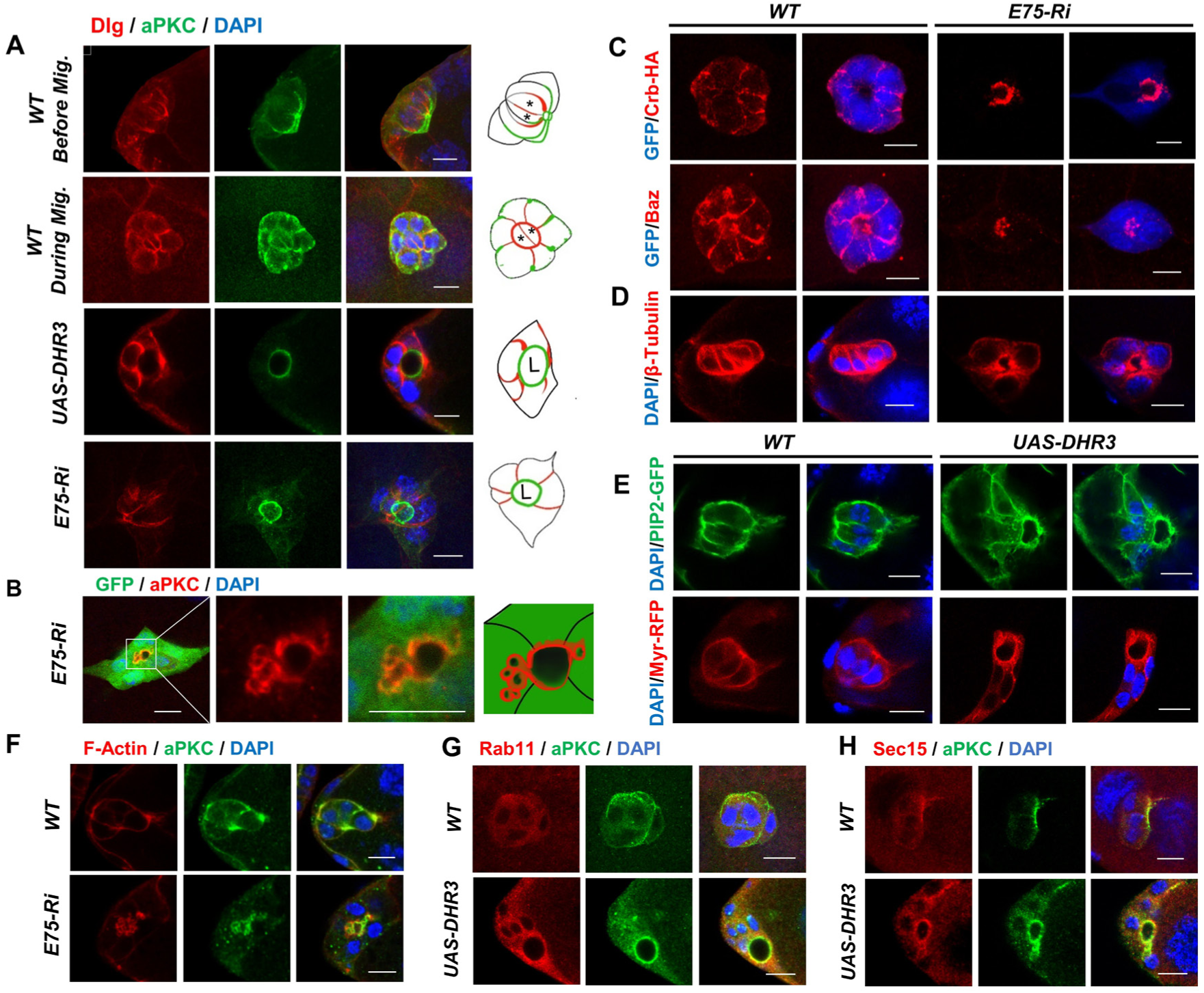
E75 loss of function and DHR3 overexpression lead to precocious lumen formation of the border cells. (A) The first two rows show confocal images of wild type border cells before migration (first row, early stage 9) and during migration (second row, mid stage 9) respectively. Before migration, the apical (stained with aPKC) and lateral (stained with Dlg) membranes of border cell cluster points to the posterior direction (to the right), with apical membrane more posterior than lateral membrane. During migration, the orientation of border cell cluster undergoes a 90 degree turn, resulting in the apical-lateral axis being perpendicular to the posterior direction (to the right). The two central polar cells are outlined by strong staining of Dlg and marked with * in the diagrams to the right. The last two rows depict border cells with *E75 RNAi* or *DHR3* overexpression that failed to migrate and instead formed lumen (marked with “L” in the diagrams) that is enclosed by aPKC stained membrane. Dlg staining is restricted to membranes between adjacent border cells. The first and last rows are resulted from maximum projection of z-stacks of confocal sections, the others are single confocal sections. (B-E) Images of border cells labeled with aPKC (B), Crb-HA and Baz (C), β-tubulin (D) staining, and PIP2-GFP and Myr-RFP (E) fluorescence, as resulted from *E75 RNAi* or *DHR3* overexpression. DAPI labels all nuclei. PIP2-GFP serves as a reporter for PIP2-enriched membrane (PLCδ-PH-GFP, see Methods for details), and Myr-RFP (myristoylated RFP) serves as a general membrane marker. (F-H) Images showing co-staining of aPKC with phalloidin (F-actin, F), Rab11 (recycling endosome marker, G), and Sec15 (exocyst component, H), as resulted from *E75 RNAi* or *DHR3* overexpression. Scale bars, 10 μm for all panels. See also Figure S3.

Formation of a tube and its enclosing lumen from non-epithelial cells is referred to as *de novo* lumen formation (Sigurbjornsdottir et al., 2014), which is a fundamental morphogenetic process central to animal development. Extensive studies in various *in vitro* and *in vivo* model systems have revealed that the initial stage of *de novo* lumen formation involves establishment of a new apical-basal polarity, which requires re-routing of multiple cellular processes and components including polarized intracellular trafficking, polarized actin and microtubule cytoskeleton, polarized distribution of apical markers, and newly synthesized membrane (Akhtar and Streuli, 2013; Datta et al., 2011; Sigurbjornsdottir et al., 2014). We found that in addition to the re-distribution of apical markers to the lumen-facing membrane, the intracellular traffic as well as cytoskeleton was also dramatically re-organized in the *E75 RNAi* or *DHR3* overexpressing border cells. Staining with Rab11 and Sec15 antibodies revealed that recycling endosome and exocyst were enriched in the cytoplasmic regions near the lumen-facing apical membrane, indicating a polarized transport toward the lumen (Figures 2G and 2H). Furthermore, F-actin and, sometimes, aPKC were observed localizing to large vacuole-like compartments adjacent to the lumen-facing membrane (Figures 2B and 2F), suggesting that these large vesicles could be in the process of fusing with the adjacent apical membrane. This phenomenon was similar to previous reports of VACs (vacuolar apical compartments) forming in the MDCK cells that are undergoing *de novo* lumen formation (Brignoni et al., 1993; Vega-Salas et al., 1988). In addition, the actin and microtubule cytoskeletons were re-organized in such a way that they are now mostly localized in and adjacent to the lumen-facing membrane. Interestingly, β-tubulin was re-organized into a distribution pattern that seems to radiate away from the central lumen (Figure 2D). Lastly, marked increase of intracellular membrane levels as indicated by Myr-RFP and PIP2-GFP was observed in the cytoplasm of *DHR3* expressing border cells, suggesting that high levels of newly synthesized membranes are needed for formation and expansion of lumen-facing membrane (Figure 2E). Taken together, these results indicate that during border cell migration E75 acts to suppress DHR3’s lumen formation function, which includes re-routing of endocytic recycling, re-distribution of apical markers, re-polarization of actin and microtubule cytoskeletons, and increased levels of membrane components.

### DHR3 is later required for the formation of micropyle tip

We next sought to understand why E75 needs to suppress DHR3’s lumen formation function during border cell migration. After border cells finished their anterior migration to the border at stage 10A, they will further migrate a short distance dorsally and finally stop at the dorsal border between nurse cells and oocyte at stage 10B. About three hours later, around stages 12 and 13, this cluster of border cells will undergo a morphogenetic transformation to form part of the micropyle, which is a tubular structure essential for sperm entry (Montell et al., 1992). Such a morphogenetic process is not well characterized and understood. Therefore, we wonder whether the lumen-forming phenotype from *E75* knockdown or DHR3 over-activation represents the precocious occurrence of the morphogenetic event involved in micropyle formation. If that is the case, E75 may be actually preventing a late morphogenetic process from occurring earlier (i.e. before or during border cell migration). Therefore, E75 and DHR3 may function together to keep the correct temporal order between the two morphogenetic processes. To address this possibility, we first sought to characterize and understand the process that enables wild type border cell cluster to be transformed into the tip of micropyle.

Collective migration of border cells has been extensively studied, but the morphogenetic process that turns the border cells into micropyle tip is little studied. Previous work by Montell and coworkers first demonstrated that border cells develop into the tip of micropyle and contribute to the cellular process thought to maintain a functional opening, while the centripetal follicle cells form the bulk of the micropyle structure. Furthermore, in the absence of border cells, a slightly smaller micropyle structure could still form, but it lacks the functional opening required for sperm entry (Montell et al., 1992). We sought to describe and characterize such morphogenetic process in details, using markers of actin cytoskeleton, membrane and apical polarity (Figures 3A-3D, Movies S2-S4). Similar to migratory border cells at stage 9, border cells at stage 10 (a period of about 10 hours, Figure 3A) mostly retain the coherent cluster morphology as well as the distribution pattern of F-actin and apical polarity proteins. During stages 9 and 10, Par6-GFP was shown to localize between adjacent border cells in a thin section of junctional region (Figure 3B, Movie S2), which was subsequently retracted and significantly shortened during stage 11 (a period of about 0.5 hour). During stage 12, Par6-GFP localization is further remodeled, with its pattern shifted from junctional region between adjacent border cells to the membrane facing the lumen-like cavity (Figure 3B, Movie S2). Consistently, Lifeact-GFP and a PIP2 membrane reporter (PIP2-GFP, Cliffe et al., 2017) both demonstrate a similar remodeling in their distribution patterns from stage 9 to stage 12, with Lifeact-GFP and PIP2-GFP highly enriched in the same membrane region enclosing the luminal space in wild type border cells at stage 12 (Figures 3C and 3D, Movies S3 and S4). A very small percentage of wild type stage 11 or 12 egg chambers would contain border cells that failed to migrate properly and reach the oocyte border (Figures S4A-S4E). Interestingly, we found that those stages 11 and 12 border cells with migration defects also displayed lumen formation that was accompanied by the remodeling of apical markers, F-actin, and PIP2-enriched membrane and was similar to the DHR3-induced lumen formation process at stages 9 and 10 (Figures S4A-S4E). This result indicates that the remodeling process is autonomously initiated in border cells and is under strict temporal control. Finally, *DHR3* knockdown or *E75* overexpression each led to disruption of the remodeling process (Figures 3E and 3F). As shown by the PIP2-GFP marker (Figure 3E), most of *DHR3 RNAi* or *E75* overexpressing border cell clusters at stages 12 and 13 displayed a cluster morphology that is characteristic of border cells at stages 9 and 10 (Figure 3C, Movie S3), where the PIP2-GFP is broadly localized in membranes between adjacent border cells. Consequently, these border cells failed to develop into the anterior tip of micropyle that surrounds a lumen-like cavity. Together, these results demonstrate that DHR3 activity is required for the morphogenetic process of lumen formation that is essential to micropyle formation. Interestingly, the morphogenetic remodeling process involved in micropyle formation is similar to the DHR3-induced lumen formation process occurred precociously in border cell cluster during stages 9 and 10, suggesting that DHR3 is not only required but also sufficient for all the remodeling events necessary for lumen formation. Indeed, random ectopic expression of *DHR3* in small clones of follicle cells by the Flip-out technique could sometimes induce formation of lumen-like structures enclosed by *DHR3* expressing follicle cells (Figure S3C), supporting the above idea.

**Figure 3.**
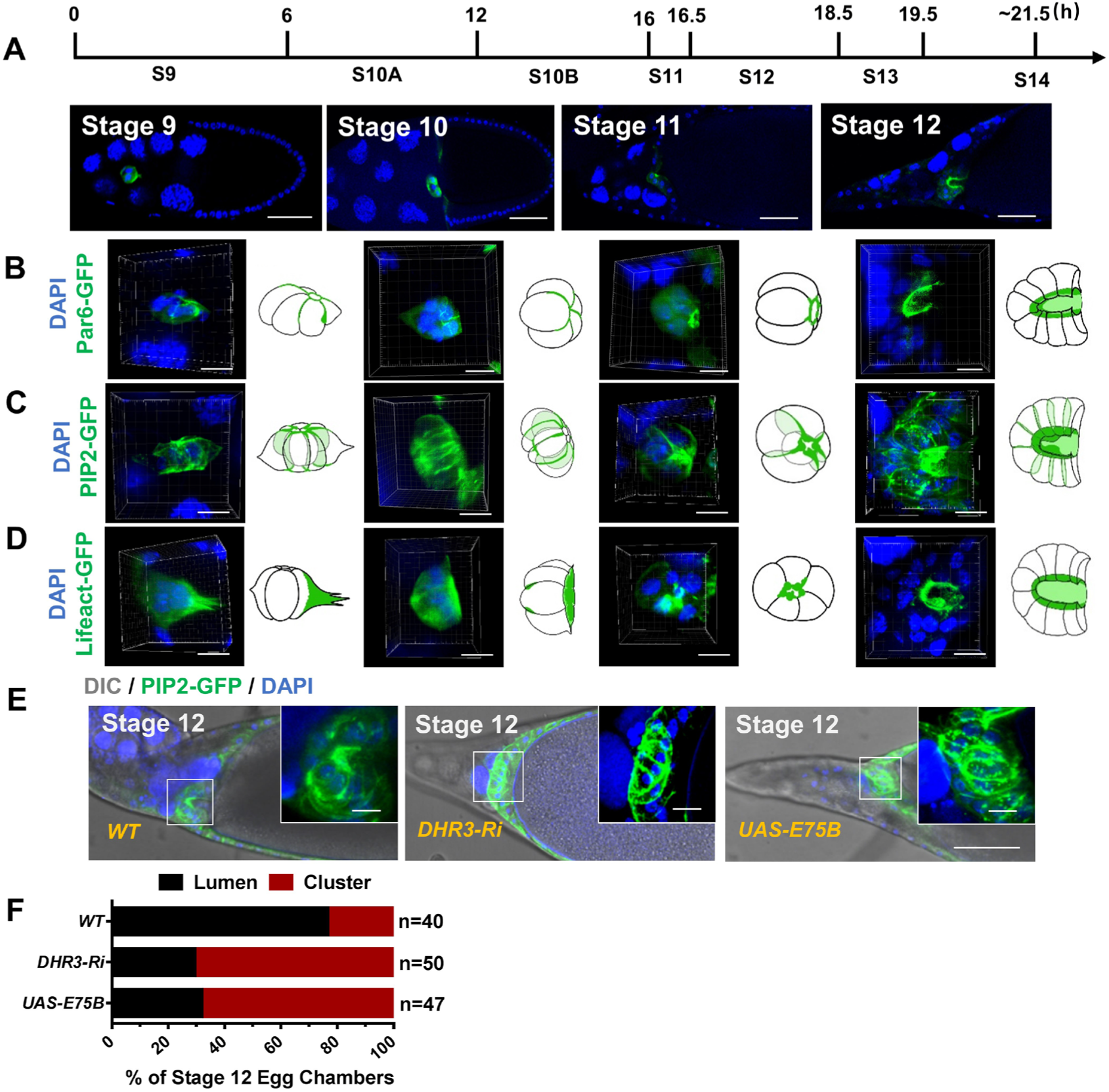
DHR3 is required for border cells’ lumen formation in the micropyle at stage 12. (A-D) A time course of developing wild type egg chambers at stages 9, 10, 11 and 12. 3-D reconstruction of z-stacks of confocal section (see Methods for details) reveals the change of morphology from a cluster to the anterior portion of the tubular micropyle (B-D). See also Movies S2-S4 that are generated from the 3-D reconstruction. Par6-GFP (B), Lifeact-GFP (C) and PIP2-GFP (D) fluorescence displays a dynamic remodeling of apical polarity, F-actin and PIP2-enriched membrane in border cells during micropyle formation. The genotype of the PIP2-GFP reporter (Cliffe et al., 2017) is detailed in Methods. (E, F) *DHR3 RNAi* and *E75B* overexpression each caused disruption of lumen formation, as compared to the morphology of wild type border cells (outlined by PIP2-GFP) at stage 12. Their cluster or lumen morphology are quantified in (F). 76.6% of stage 12 wild type border cell clusters displayed obvious lumen morphology, whereas 70.4% of *DHR3 RNAi* and 68.0%, of *E75B* overexpression displayed cluster morphology, which is characteristic of the wild type border cells at stage 10 (C). Scale bars, 50 μm in (A, E), 10 μm in high-magnification views in (B-E). See also Figures S3 and S4.

### Reduction of EcR signaling and E75 levels causes de-repression of DHR3 activity

We next sought to understand how DHR3 function is temporally regulated to limit its lumen forming activity only to the period of micropyle formation and not to the period of collective migration. We reasoned that DHR3’s activity in border cells has to be inhibited by E75 during stages 9 and 10, as shown by our results above (Figure 1). Afterward, DHR3’s activity would need to be de-repressed beginning at stage 11 to start the morphogenetic process of lumen formation. We already showed that DHR3 function is antagonized by E75, and that both E75 and DHR3 are expressed by EcR during border cell migration at stage 9. We then examined the temporal expression patterns of E75 and DHR3 as well as the levels of EcR signaling. We found that EcR signaling, as reflected by its well-established reporter *EcRE-lacZ*, reached its highest levels during stage 10B, and then dramatically declined from stage 11 to stage 13 (Figures 1C, 4A and 4E). Accordingly, both expression levels of the *E75-lacZ* reporter, which reflects the transcription levels of *E75* (Figures 4B and 4F), and the protein levels of DHR3 as detected by DHR3 antibody also decreased from stage 11 to stage 13 (Figures 4C and 4G). These results suggest that as ecdysone signaling decrease dramatically (beginning at stage 11) E75 level should also decrease to a low level (at stages 11 and 12), which may be below the threshold level for inhibition of DHR3’s activity. To test this possibility, we need a good activity reporter for DHR3’s function.

**Figure 4.**
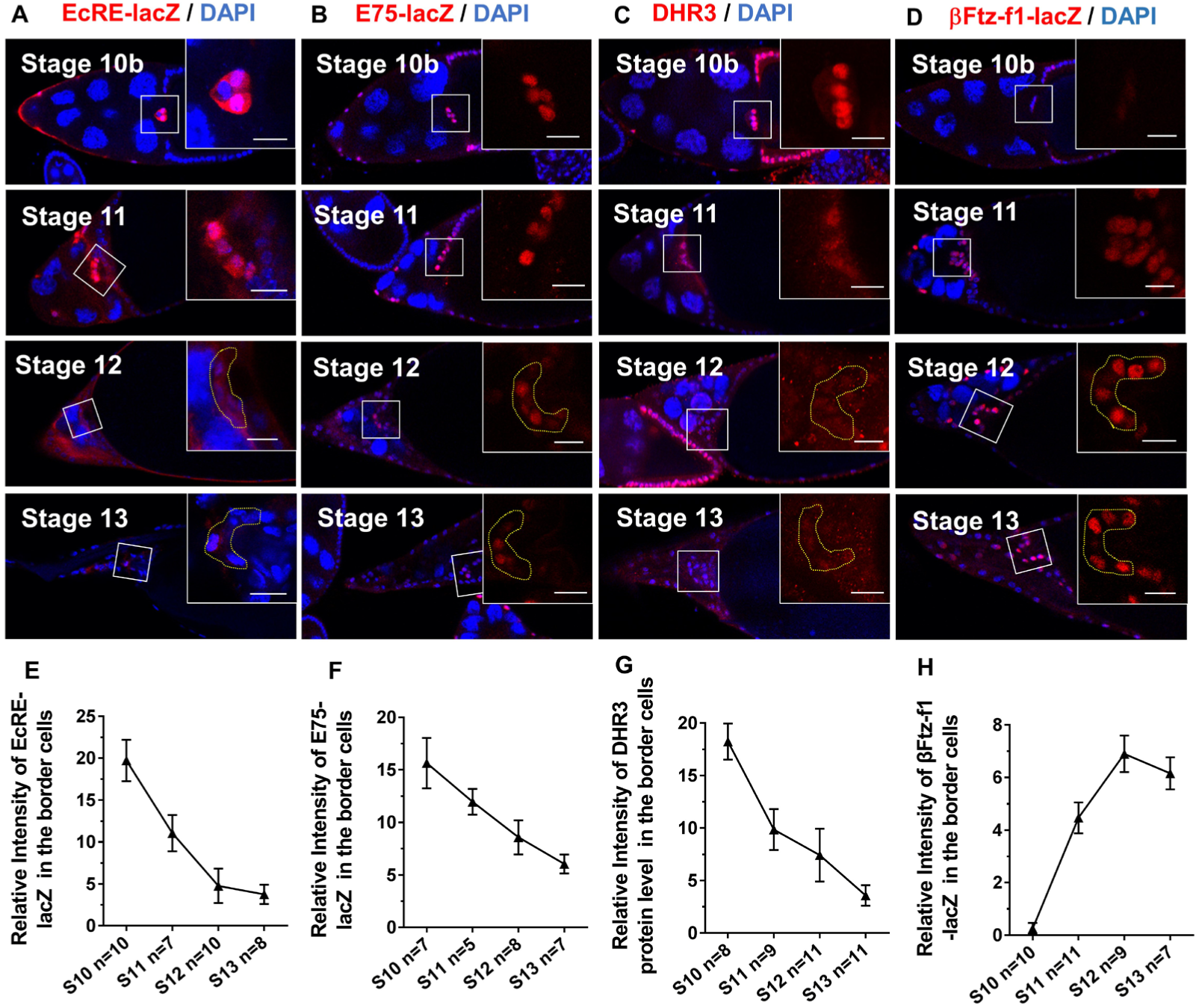
Temporal expression patterns of EcRE-lacZ, E75-lacZ, DHR3 and β-Ftz-f1-lacZ from stage 10b to stage 13. (A-D) Confocal images showing antibody staining of β-gal that is expressed by the *EcRE-lacZ* reporter (A), *E75-lacZ* enhancer trap (B), and *β-Ftz-f1-lacZ* enhancer trap (D), as well as antibody staining of DHR3 (C), from stage 10b to stage 13. Boxed regions are enlarged and shown at the right of all panels. Areas encircled by yellow dotted lines (based on labeling of GFP as expressed by *slbo-Gal4*) highlight the border cell clusters (A-D) at stages 12 and 13. Scale bars, 10 μm. (E-F) Quantification of antibody staining of border cell clusters in (A-D) from stage 10b to stage 13. The number of egg chambers examined (n) for each stage is given at the x-axis. Statistical analysis was performed using two-tailed Student’s *t*-test. Error bars indicate S.E.M. See also Figure S5.

Previous literatures indicate that DHR3’s immediate downstream target gene during metamorphosis is *βFtz-f1* (Geanette T. Lam1, 1997; Jia et al., 2017; Kageyama et al., 1997), whose expression levels serve as a readout for DHR3 activity. We obtained an enhancer trap line for *βFtz-f1, βFtz-f1-lacZ*, which has a *lacZ* containing P-element inserted in the 5’ UTR region of the gene and supposedly could reflect the transcription level of *βFtz-f1*. We found that its expression could serve as a bona fide reporter for DHR3 activity, based on the following results. First, *βFtz-f1-lacZ* expression is initially at non-detectable levels at stages 9 and 10A (Figure S5A), and at very low levels at stage 10B (Figures 4D and 4H), then it abruptly reaches much higher levels at stages 11, 12 and 13 (Figure 4D,H). Therefore, *βFtz-f1-lacZ*’s temporal expression pattern is highly consistent with our above prediction about the temporal regulation of DHR3 activity. Second, *DHR3* overexpression led to precocious expression of *βFtz-f1-lacZ* within border cells during stages 9 and 10, whereas co-expression of *E75B* and *DHR3* suppressed such precocious expression (Figures 5A and 5C). Conversely, *DHR3 RNAi, E75B* overexpression, or *E75B* and *DHR3* co-expression, each inhibited *βFtz-f1-lacZ*’s normal expression in border cells during stage 11 (Figures 5B and 5D). Furthermore, *DHR3* overexpression in the follicle cells at the stage 9, when *βFtz-f1-lacZ* is not normally expressed, ectopically induced *βFtz-f1-lacZ*’s expression in the follicle cells (Figure S5B). Third, expression of *βFtz-f1-RNAi* in the background of *E75 RNAi* partially rescues border cell’s migration defects (Figure 5E and Figure S6A), suggesting that βFtz-f1 functions downstream of DHR3. Lastly, expressing *βFtz-f1-RNAi* in the border cells resulted in the disruption in the formation of micropyle tip, similar to the loss-of-function defects of *DHR3 RNAi* (Figures S6B and S6C). On the other hand, overexpression of *βFtz-f1* in stage 10 border cells resulted in actin-enriched patches that are similar to PAPs from moderate *E75 RNAi* defects (Figures S3A and S3B), suggesting an incomplete lumen forming phenotype. Taken together, these results support the conclusion that reduction in EcR signaling and E75 levels leads to de-repression of DHR3 activity (as represented by the *βFtz-f1-lacZ* reporter) beginning at stage 11, which serves to switch on lumen formation for micropyle formation (during stages 11 to 13). Moreover, βFtz-f1 acts downstream of DHR3 to mediate micropyle formation.

**Figure 5.**
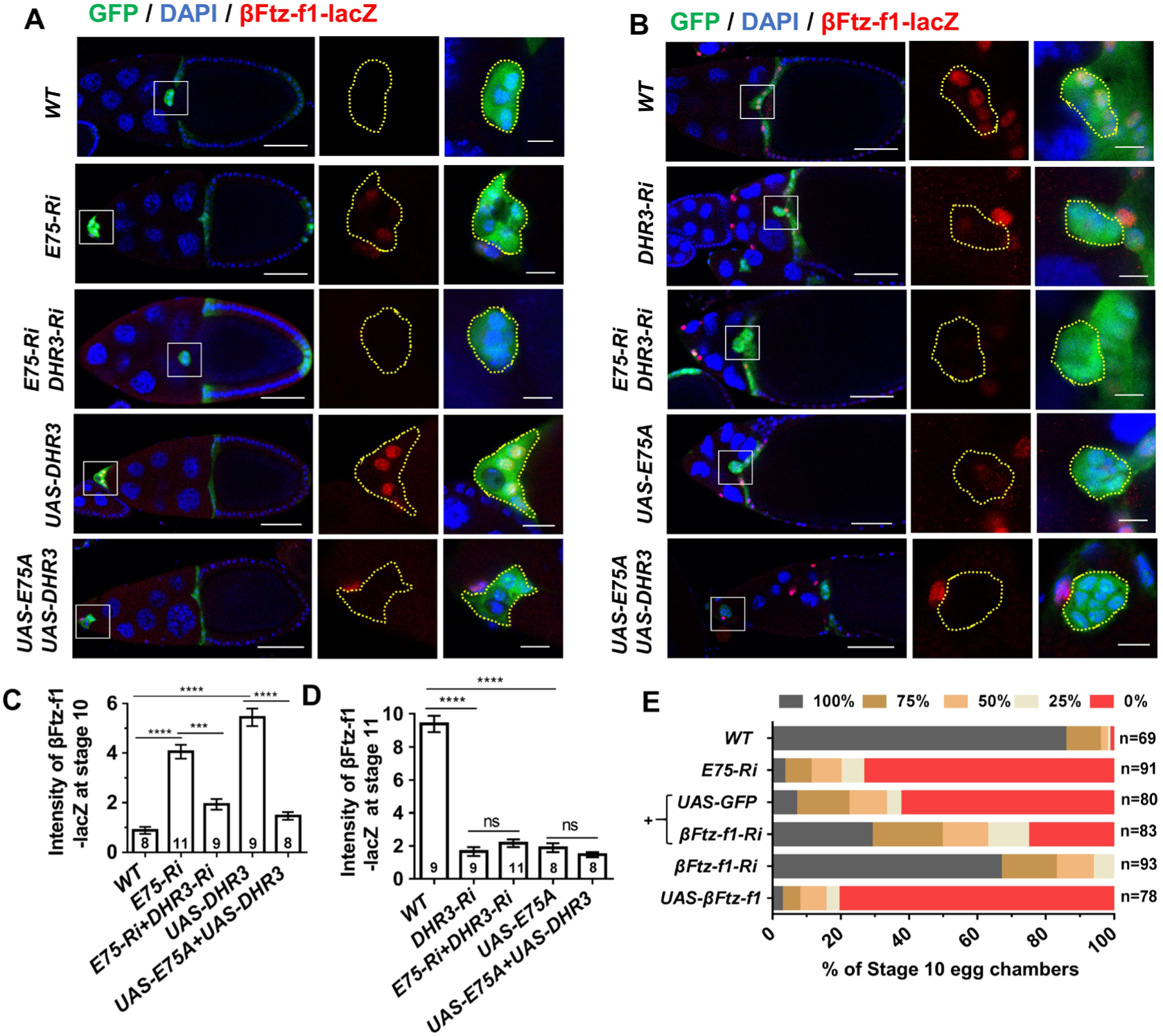
β-Ftz-f1 acts downstream of DHR3 and its expression serves as a reporter of DHR3 activity. (A) β-Ftz-f1-lacZ levels in border cells at stage 10 as represented by β-gal antibody staining. Compared to wild type (WT) control, *E75 RNAi* and *DHR3* overexpression both resulted in significant increase of β-Ftz-f1-lacZ levels (quantified in C), while double knock down of *E75* and *DHR3* (*E75 Ri + DHR3 Ri*) and overexpression of both *DHR3* and *E75* (*UAS-DHR3 + UAS-E75*) abolished the increase (quantified in C). (B) β-Ftz-f1-lacZ levels in border cells at stage 11 as represented by β-gal staining. Compared to wild type (WT) control, *DHR3 RNAi* and *E75* overexpression both resulted in significant reduction of β-Ftz-f1-lacZ levels (quantified in D). Yellowed dotted lines (A, B) outline individual border cell clusters, as labeled with GFP expressed by *Slbo-Gal4*. Boxed regions (A, B) are enlarged and shown at the right of all panels. Scale bars, 50 μm for egg chambers, 10 μm for border cells. (C, D) Quantification of β-Ftz-f1-lacZ levels. The number of egg chambers examined for each genotype is indicated within its corresponding column. Statistical analysis was performed using two-tailed Student’s *t*-test. Error bars indicate S.E.M. **, P<0.01; ***, P<0.001; ****, P<0.0001; ns, not significant. (E) Quantification of rescue of border cell migration defects of *E75 RNAi* by co-expression of *β-Ftz-f1 RNAi*. Represented images for the indicated genotypes are shown in Figure S6A. See also Figure S6.

### DHR3 is required and sufficient for chitin secretion into the lumen

An essential feature of *de novo* lumen formation in the vertebrates is the secretion of glycoprotein such as the negatively charged podocalyxin into the lumen to keep the lumen membranes apart and promote the expansion of luminal space (Bryant et al., 2014; Strilic et al., 2010). Although *Drosophila* does not possess a podocalyxin homolog, the tube formation during *Drosophila* tracheal development requires the secretion of chitin into the lumen (Devine et al., 2005). Chitin is a long-chain polymer of N-acetylglucosamine, which is also a primary component of the *Drosophila* exoskeleton (Moussian et al., 2005; Zhu et al., 2016). We then proceeded to determine whether chitin is present in the lumen enclosed by the border cells and whether DHR3 acts to promote secretion of chitin into the lumen. Interestingly, we found that chitin (labeled by FB-28) is only present in the extracellular space adjacent to wild type border cells during and after stage 11 (Figures 6B, 6D, 6E and S7E), whereas it is not present around the border cells before stage 11 (Figures 6A, 6C and S7E). A very small percentage of wild type stage 12 egg chambers would contain border cells that failed to migrate properly and reach the oocyte border, and we found that chitin is present within the lumen surrounded by those border cells (Figure 6F). Together, these results indicate that chitin is present in the lumen within the border cell cluster. In addition, the temporal and localization patterns of chitin suggest that it is secreted by border cells beginning at stage 11 during lumen formation for the micropyle tip. Furthermore, we found that expressing *E75 RNAi* or *DHR3* specifically in the border cells (by *slbo-Gal4*) each resulted in chitin being precociously localized within the lumen of border cell clusters that failed to migrate to the oocyte border at stage 10 (Figures 6A and 6C), indicating that DHR3 activation is sufficient to induce chitin secretion. On the contrary, *DHR3* knockdown or *E75* overexpression led to loss of extracellular chitin near border cells at stage 11 (Figures 6B and 6D), indicating DHR3 is required for chitin secretion by the border cells. Together, these results indicate that DHR3 activity is necessary and sufficient for chitin secretion by the border cells during lumen formation. To further test whether βFtz-f1 is also sufficient for chitin secretin, we examined and found no chitin secretion in *βFtz-f1* overexpressing border cells at stage 10 (Figures S7A and S7C). This result could be due to the aforementioned fact that *βFtz-f1* overexpression only resulted in PAP (incomplete lumen formation, Figure S6A). Hence, it is conceivable that chitin secretion can only occur after lumen formation progresses to a certain degree. On the other hand, we found that βFtz-f1 is required for chitin secretion by the border cells during micropyle formation (Figures S7B and S7D), similar to DHR3’s role (Figures 6B and 6D). These results suggest that βFtz-f1 may act downstream of DHR3 to partially mediate DHR3’s chitin secretion role.

**Figure 6.**
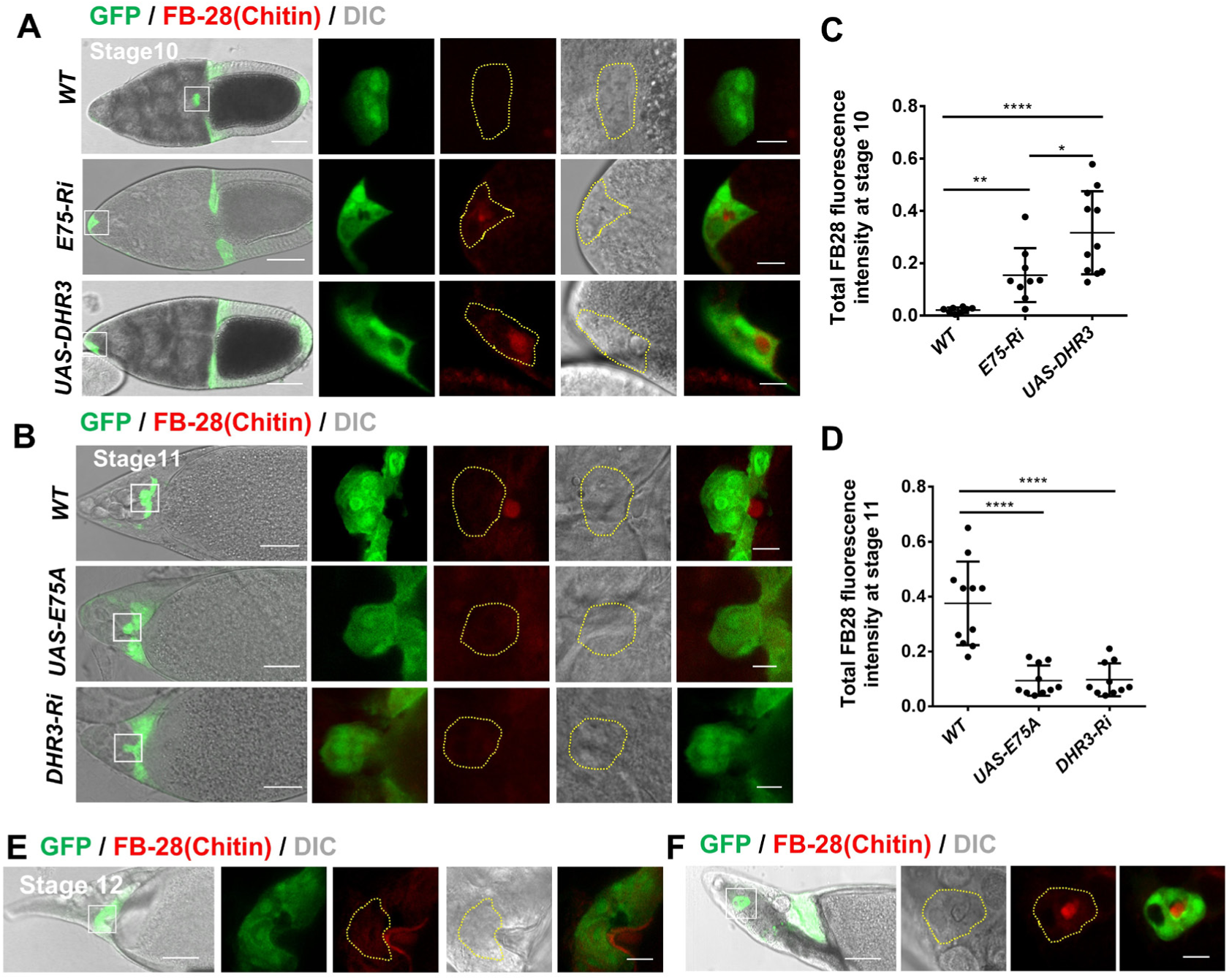
DHR3 is necessary and sufficient for chitin secretion by the border cells during lumen formation. (A, B) Confocal and DIC images showing chitin staining in stage 10 (A) and stage 11 (B) egg chambers. Chitin is labeled with the Fluorescent Brightener 28 (FB28) dye. (A) Chitin was not detected within or adjacent to wild type (WT) border cells at stage 10, whereas *E75* RNAi and *DHR3* overexpression in the border cells resulted in precocious secretion of chitin to the lumen (quantified in C). (B) Starting at stage 11, chitin was detectable adjacent to WT border cell cluster, but *DHR3* RNAi or *E75* overexpression abolished chitin staining (quantified in D). Yellow dotted lines outline individual border cell clusters as labeled with GFP expressed by *Slbo-Gal4*. Scale bars, 50 μm for egg chambers, 10 μm for border cells. (C, D) Quantification of chitin levels of border cells for the indicated genotypes. (E) At stage 12, chitin was detectable in the lumen of micropyle, where wild type border cells had completed their migration. (F) In stage 12 wild type border cells that exhibited migration defects, chitin was detected in the lumen enclosed by border cells. Statistical analysis was performed using two-tailed Student’s *t*-test. Error bars indicate S.E.M. **, P<0.01; ****, P<0.0001. See also Figure S7.

### DHR3 and βFtz-f1 suppress JNK signaling in the border cells

Lastly, we sought to explore what signaling pathways DHR3 regulates in border cells. We tested reporters for a number of signaling pathways previously known to play essential roles in the border cells, including JAK/STAT (Beccari et al., 2002; Silver et al., 2005), Notch (Wang et al., 2007), JNK (c-Jun N-terminal kinase) (Llense, 2008; Melani et al., 2008) and Dpp (Luo et al., 2015). Among them, JNK was the only signaling found to be severely affected by E75 knockdown or DHR3 overexpression (Figure S8A). JNK signaling pathway was previously reported to be required for cell-cell adhesion between adjacent border cells during their collective migration (Llense, 2008; Melani et al., 2008). Staining for *Puc-lacZ*, a widely used reporter for JNK signaling, revealed that both *E75 RNAi* and *DHR3* overexpression caused strong reduction of *Puc-lacZ* reporter activity in stage 10 (Figures 7A-7C), indicating that increased DHR3 activity suppresses JNK signaling. Indeed, knockdown of *DHR3* in the background of *E75 RNAi* rescued the level of *Puc-lacZ* expression back to the wild type level in stage 10 (Figures 7A and 7B). Furthermore, overexpressing *bsk* (encoding *Drosophila* JNK) in the background of *E75 RNAi* or *DHR3* overexpression partially rescued the severe migration defects and precocious lumen formation of border cells that were resulted from *E75* loss of function (Figure 7D and S8B). Together, these results demonstrate that reduction of E75 or increase of DHR3 activity leads to downregulation of JNK signaling in the migratory border cells at stages 9 and 10. We next tested whether JNK signaling was negatively regulated by DHR3 and βFtz-f1 during micropyle formation. We showed that in the wild type the level of JNK signaling was reduced from stage 10 to stage 11, and then further reduced from stage 11 to stage 12 (Figures 7E and 7F). In stage 12 border cells, *DHR3* knockdown, *E75* overexpression, and *βFtz-f1* knockdown each increased the originally low JNK signaling to a much higher level, which is similar to the level at stage 10 (Figures 7E and 7F). These results suggest that JNK signaling needs to be suppressed in order for lumen formation to occur properly during stages 11 and 12. Indeed, overexpression of *bsk* and hence increase of JNK signaling resulted in disruption of lumen formation during formation of the micropyle tip at stage 12 (Figures 7G and 7H). Taken together, these results suggest that JNK-mediated cell adhesion between border cells is temporally and differentially regulated during two different morphogenetic processes: collective migration and micropyle formation, and that its downregulation by DHR3 and βFtz-f1 is essential for lumen formation in the latter process.

**Figure 7.**
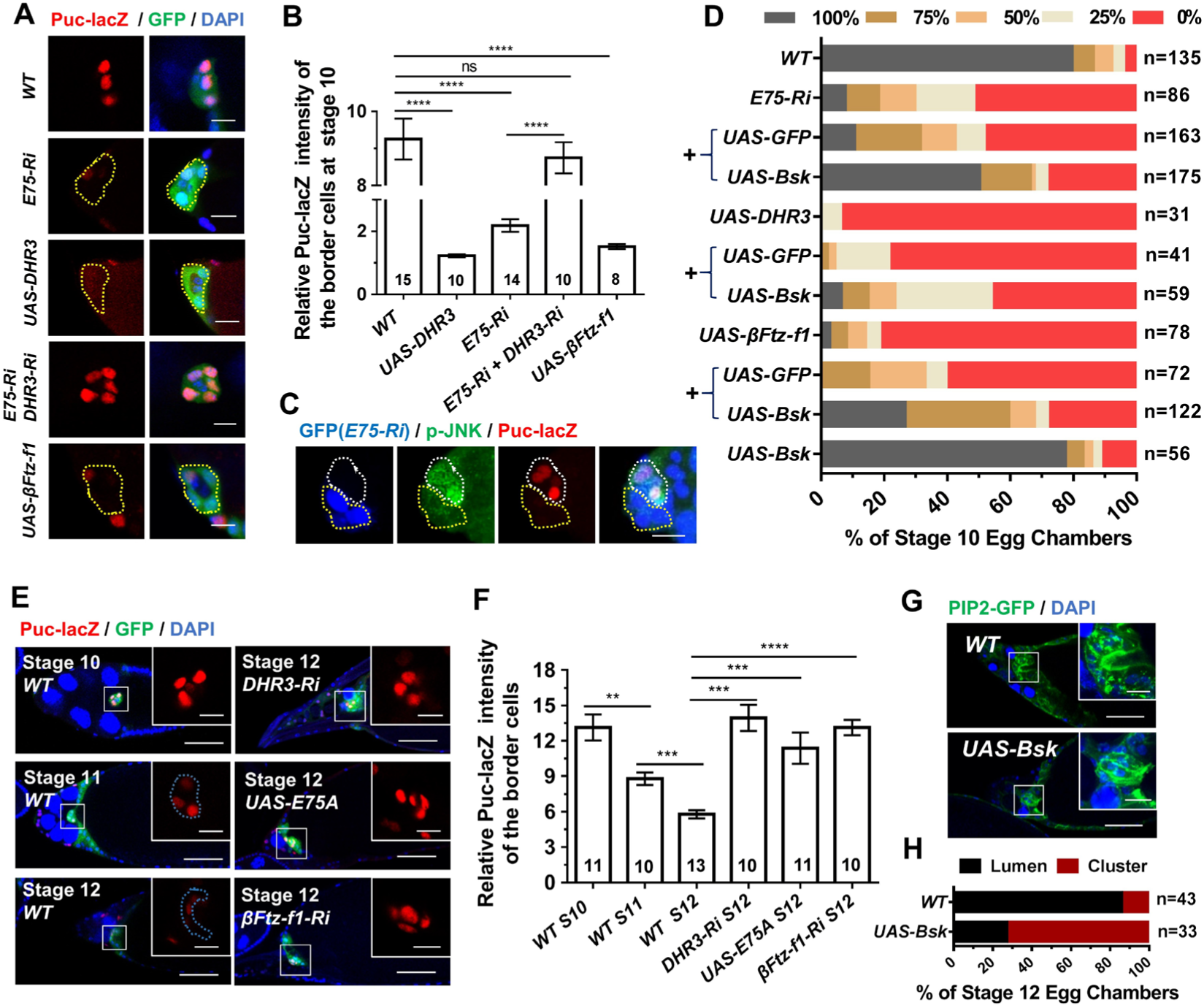
DHR3 and βFTZ-f1 downregulate JNK signaling in the border cells. (A, B) *Puc-lacZ* expression levels in migratory border cells at stage 9 or 10 as represented by β-gal antibody staining. *E75 RNAi, DHR3* and *βFTZ-f1* overexpression each resulted in strong and significant decrease of *Puc-lacZ* levels as compared to wild type (WT) control (quantified in B), while coexpression of *DHR3 RNAi* and *E75 RNAi* returns the *Puc-lacZ* levels to that of WT (quantified in B). (C) A mosaic border cell cluster containing a clone of *E75 RNAi* expressing cells (marked by GFP, outlined with yellow dotted line), which exhibited reduction of *Puc-lacZ* and p-JNK levels as compared to those in the adjacent wild type cells (no GFP, outlined with white dotted line). (D) Quantification of partial rescue of border cell migration defects of *E75 RNAi, DHR3* overexpression, and *βFTZ-f1* overexpression by coexpression of *bsk*. (E, F) *Puc-lacZ* expression levels in WT border cells decreased from stage 10 to stage 12, while expression of *DHR3 RNAi, E75A* and *βFTZ-f1 RNAi* elevated *Puc-lacZ* levels in border cells at stage 12. The results are quantified in (F). (G, H) *bsk* overexpression caused disruption of lumen formation, as compared to morphology of wild type border cells (outlined by PIP2-GFP) at stage 12. Their cluster or lumen morphology are quantified in (H). 86.0% of stage 12 wild type border cells displayed obvious lumen morphology, whereas 72.7% of *bsk* overexpressing cells displayed cluster morphology, which is characteristic of the wild type border cells at stage 10. Statistical analysis was performed using unpaired two-tailed Student’s *t*-test. Error bars indicate S.E.M. **, P<0.01; ***, P<0.001; ****, P<0.0001; ns, not significant. Scale bars, 10 μm. See also Figure S8.

## Discussion

We demonstrate that two nuclear receptors, E75 and DHR3, are critical for temporal coordination of two very different morphogenetic processes of the border cell cluster, namely its collective migration and its lumen formation. First, our results revealed that the levels of E75 and DHR3 (in response to ecdysone) are the underlying control of the temporal order (Figure 8). Strong loss of function of *E75* or *DHR3* overexpression disrupts the temporal order and causes lumen formation to occur first. Consequently, collective migration could not take place afterward, because of the unique nature of the lumen structure, which precludes migration from occurring. Second, levels of E75 and DHR3 together with the antagonism between the two nuclear receptors underlie the mechanistic control of time interval between the two morphogenetic processes (Figure 8). E75 acts as a molecular timer. Its expression level determines the length of interval between migration and lumen formation (Figure 8). Very little E75 (strong loss-of-function) causes lumen formation to occur before migration could take place, effectively resulting in no interval between the two morphogenetic processes. Moderate *E75* loss-of-function phenotype demonstrates that collective migration could take place at early stage 9 (Figures S3A and S3B), but accompanied with a precocious occurrence of lumen formation at late stage 9 or stage 10, indicating a shortened interval. On the other hand, too much E75 (*E75* overexpression) results in reduced occurrence of lumen formation at stage 12 or 13 (Figure 3F), suggesting an expanded interval. Finally, it is important to note that delayed wild type border cells that supposedly contain the wild type levels of E75 and DHR3 exhibit a normal time interval (Figures S4 and 7F). During tissue or organ formation, it is not uncommon for a certain cell population to undergo two vastly different morphogenetic processes. This study provides a novel mechanistic insight into the molecular machinery that coordinates both the order and time interval between morphological processes.

**Figure 8.**
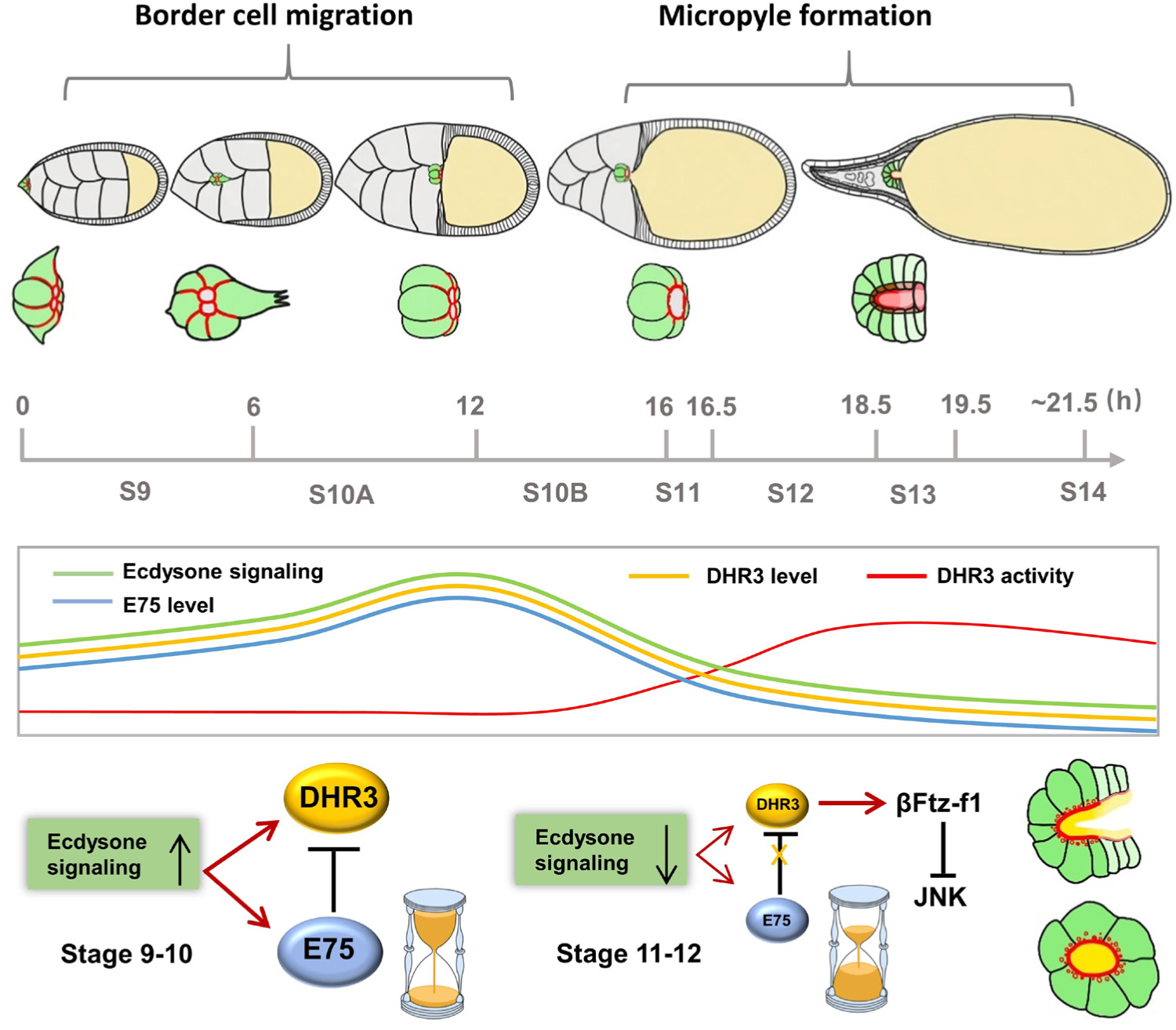
Model of how E75 and DHR3 temporally coordinate the migration and lumen formation of border cells. See description in the Discussion section for details.

Our study also uncovers a surprising mechanism of how a nuclear receptor controls the process of de novo lumen formation. DHR3 seems to act as a potent switch or inducer for lumen formation since it is necessary and sufficient for lumen formation of border cells both during stage 9 and during stages 11-13. Activation of DHR3 function in border cells seems to simultaneously induce multiple cellular processes that were previously demonstrated to be essential for *de novo* lumen formation in other systems (Sigurbjornsdottir et al., 2014), including re-routing of endocytic recycling, re-distribution of apical markers, re-polarization of actin and microtubule cytoskeletons, and increased synthesis of membrane components. In addition, DHR3 is necessary and sufficient for the secretion of chitin into the lumen of border cells both at stage 9 and at stage 12. Chitin had been previously shown to be required for tube expansion and maturation during *Drosophila* tracheal morphogenesis (Devine et al., 2005). Its function seems to provide an extracellular matrix support (Moussian et al., 2006; Wang et al., 2006). The mechanism by which chitin affects tube morphogenesis remains poorly understood. How DHR3 induces chitin synthesis and secretion and whether chitin is required for lumen formation and tube maturation in micropyle remain to be further determined. Furthermore, we demonstrate that DHR3’s lumen-inducing function is mainly mediated through βFtz-f1, a nuclear receptor and transcription factor that has been well established to be DHR3’s immediate target gene during metamorphosis. However, βFtz-f1 does not seem to mediate all of DHR3’s functions since *βFtz-f1* overexpression could not induce complete lumen structure and chitin secretion, suggesting that other factors downstream of DHR3 may also contribute to lumen formation. Lastly, we show that JNK signaling is downregulated by DHR3 and βFtz-f1, suggesting that cell adhesion between adjacent border cells needs to be reduced during lumen formation. This is consistent with the idea that remodeling of apical polarity, cytoskeleton and membrane during lumen formation may require down-regulation of cell-cell adhesion. Given the multiple functions as demonstrated for DHR3, it will be interesting to test whether these lumen inducing functions will be conserved in other developmental contexts in *Drosophila* and vertebrate. Interestingly, previous studies reported that the mammalian homolog of DHR3, RORα was enriched in human mammary duct, and its inactivation impaired polarized acinar morphogenesis (Xiong et al., 2012; Xiong and Xu, 2014), suggesting a similar role in vertebrate.

Although treated as an excellent model system for collective migration, border cells’ physiological function during oogenesis is to make a functional opening within the micropyle for sperm entry. How the border cell cluster develops into the anterior tip of the tubular structure of micropyle is poorly understood. Our study reveals a dynamic remodeling of apical polarity molecules, F-actin, and PIP2-enriched membrane, which is consistent with the process of *de novo* lumen formation. The functional roles of DHR3, βFtz-f1, EcR, E75 and JNK during micropyle formation, as demonstrated by our study, provide the first detailed analysis of this morphogenetic process. We suggest that in addition to collective migration, border cells could also serve as a model system to study *de novo* lumen formation in *Drosophila*.

## Author Contributions

Conceptualization, J.C., X.W., S.L. and G.E; Methodology, X.W., J.C., and H.W.; Investigation, X.W., H.W., L.L; Resources, S.L. and G.E.; Visualization, X.W. and H.W.; Writing, J.C., X.W. and G.E., Supervision, J.C. and H.W.

## Acknowledgments

We thank Henry Krause and Oren Schuldiner for E75 related flies and antibodies, Carl Thummel for DHR3 antibody. We also thank Jacques Montagne, Juan Huang, Lei Xue, Xiaobo Wang, Zizhang Zhou, the Bloomington Drosophila Stock Center, Tsinghua University Fly Stock Center, National Institute of Genetics Stock Center (Japan), and Vienna Drosophila RNAi Center for other fly stocks. We thank Zhenji Gan for critical comments.

## Methods and Materials

### Fly stocks

Flies were cultured and maintained on standard cornmeal media with sugar and yeast at 25°C. Progenies of crosses between *UAS-RNAi* or *UAS-transgenes* and *Slbo-Gal4* were cultured at 29°C for two days for specific gene’s knockdown and overexpression. For moderate knockdown or overexpression, flies were cultured at 29°C for only one day or cultured at 25°C, as indicated in the figure legends. To generate flip-out clones of border cells and follicle cells, female flies were heated-shocked for 3 minutes at 37°C and then kept at 29°C for 1 day before dissection.

Fly stocks listed below were obtained from different labs and stock centers, including Bloomington Stock Center (BDSC), National Institute of Genetics Stock Center, Japan (NIG), Vienna Drosophila RNAi Center (VDRC) and Tsinghua Fly Center (THFC).

### Fly stocks

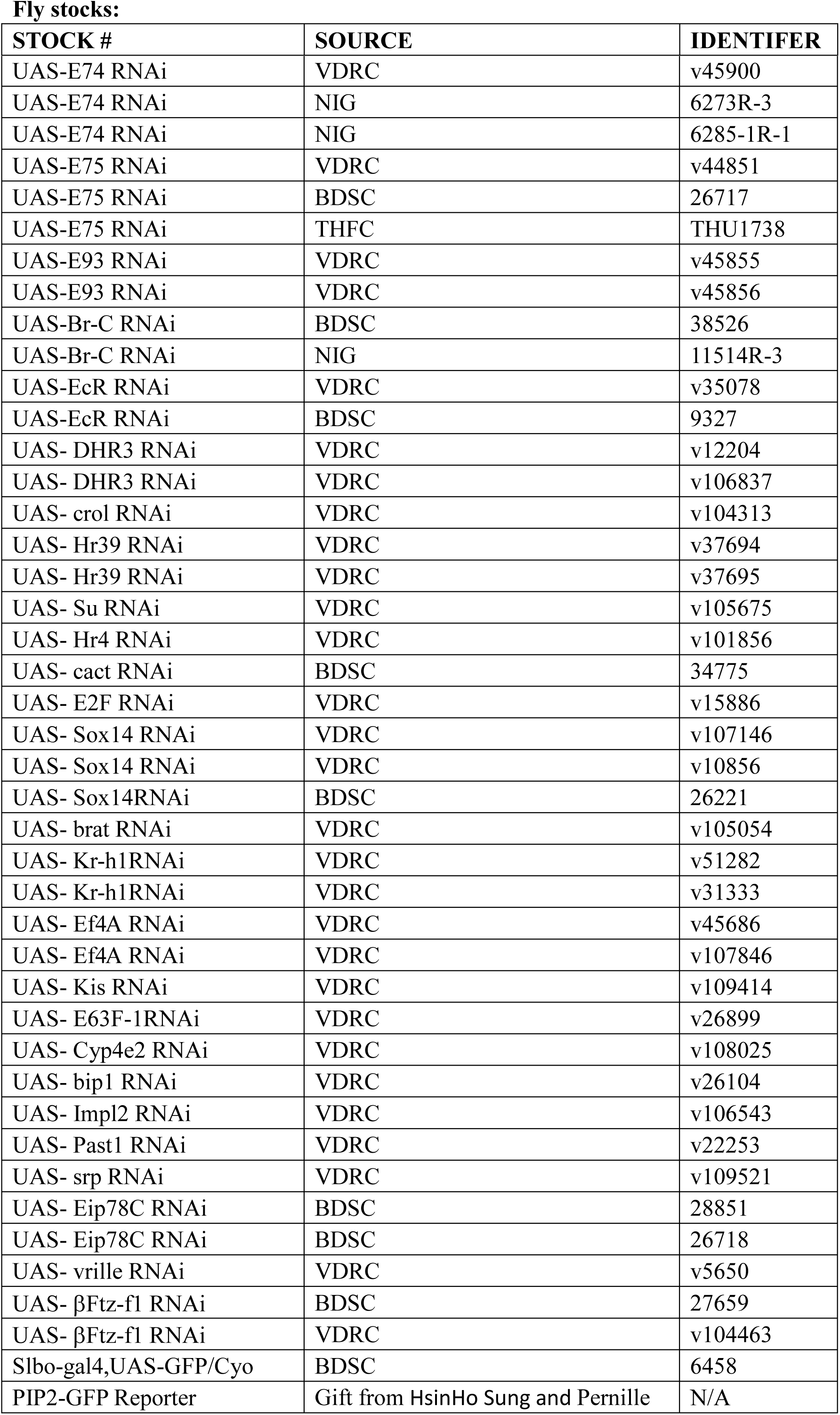
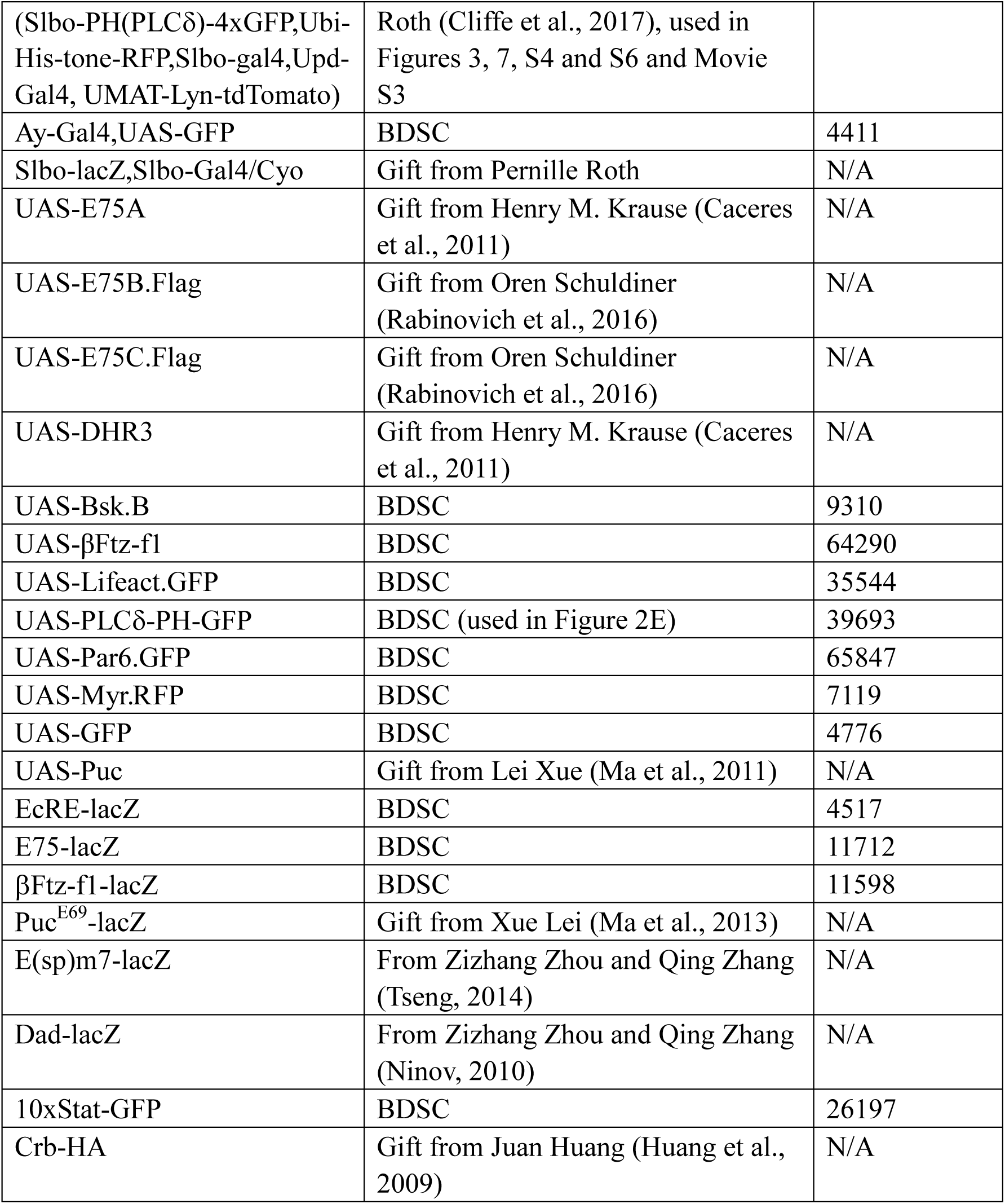

### Antibodies

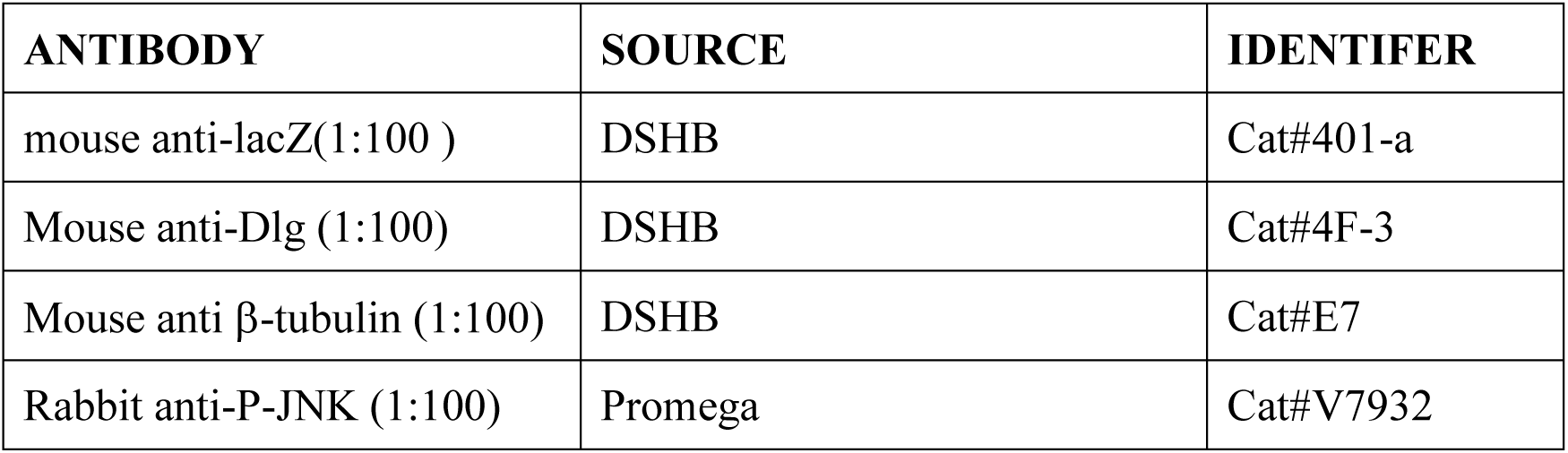
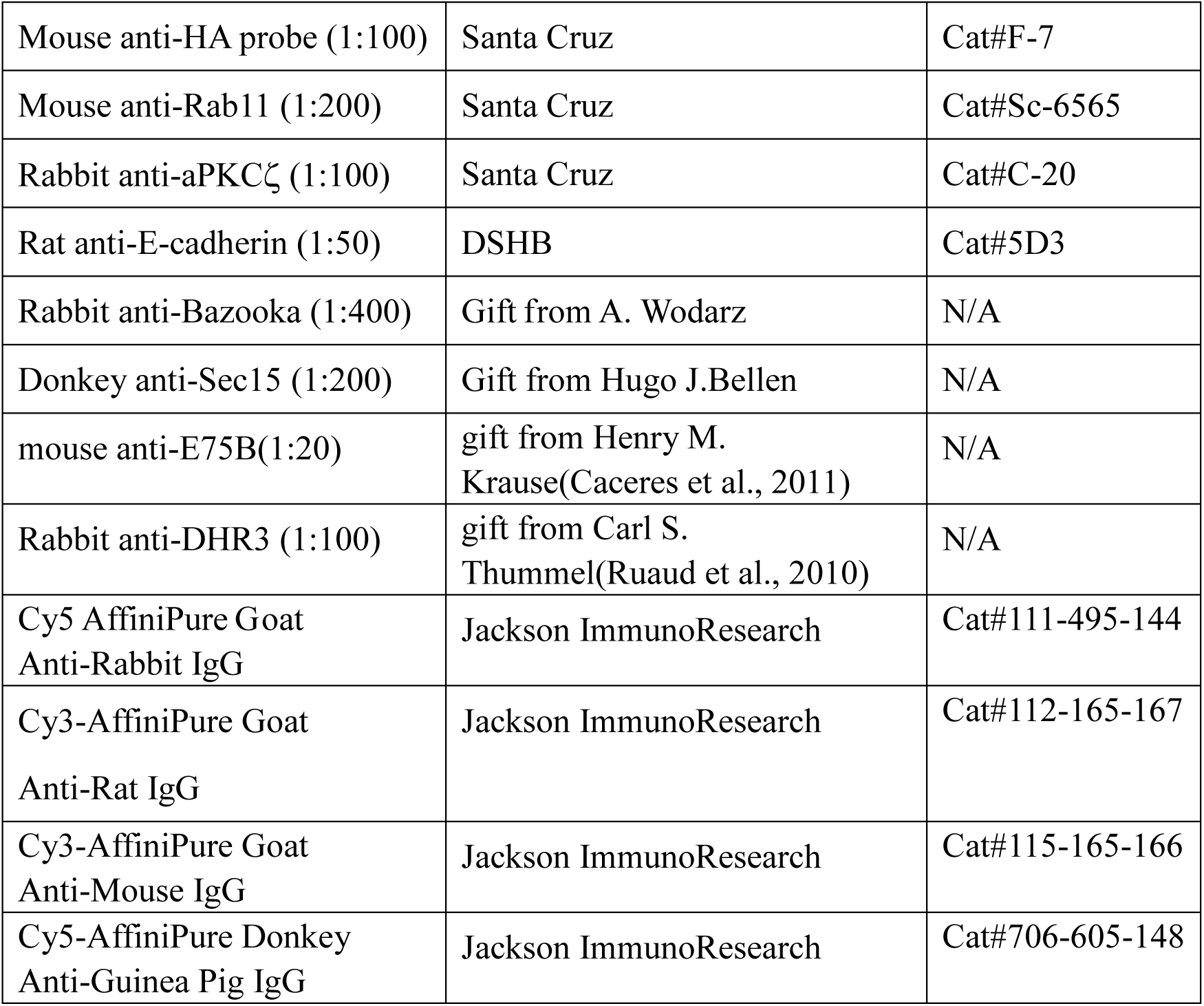

### Chemicals

| CHEMICAL | SOURCE | IDENTIFER |
| --- | --- | --- |
| Regular insulin | Novo Nordisk | NDC# 0169-2313-21 |
| Fluorescent Brightener 28 | Sigma-Aldrich | Cat#F3543 |
| DAPI | Santa Cruz Biotechnology | Cat#sc-3598; |
| TRITC-conjugated Phalloidin | Sigma-Aldrich | Cat#P1951 |

### Software

| SOFTWARE | SOURCE | IDENTIFER |
| --- | --- | --- |
| GraphPad Prism 6 | www.graphpad.com | N/A |
| Image J | http://imagej.nih.gov/ij | N/A |
| Leica confocal software | http://softadvice.informer.com/Leica_Confocal_Software.html | N/A |
| Imaris 7.2.3 | http://www.bitplane.com | N/A |

### Method Details

#### Immunostaining

Female flies were raised on fresh food with yeast at 29°C for 2 days. Ovaries were dissected in PBS, and then fixed in 100ul devitellinizing buffer (7% formaldehyde) and 600μl heptane, with strong shaking for 10 min, then washed 3 x10min with PBS, and 3×10 min with PBST. For egg chamber staining, egg chambers were blocked with 10% goat serum in PBST for 30 min after fixed and washed, and then incubated with primary antibody at 4°C overnight. Ovary samples were then washed 3×10 min with PBST, blocked with 10% goat serum in PBST for 30min, incubated with secondary antibody at 1:200 in PBST for 2 hours. DAPI was added and stained for 30 min during secondary staining. Lastly, ovaries were washed again with 10 min PBST for three times, mounted on microscope slide with 40% glycerol. Primary antibodies used include mouse anti-lacZ (1:100,401-a, DSHB), mouse anti-E75B (1:20, gift from Henry M. Krause)(Caceres et al., 2011), rabbit anti-DHR3 (1:100, gift from Carl S. Thummel)(Ruaud et al., 2010), Rabbit anti-p-JNK (1:50, Promega, V7932), Rat anti-E-cad (1:50, 5D3, DSHB), Rabbit anti-PKCζ (C-20, 1:100, Santa Cruz), mouse anti-Dlg (4F3, 1:100, DSHB), mouse anti-HA(1:100, F-7, Santa Cruz), rabbit anti-Baz (1:400, gift from A. Wodarz). Secondary antibodies were used including Cy5 AffiniPure Goat Anti-Rabbit IgG, Cy3-AffiniPure Goat Anti-Rat IgG, Cy3-AffiniPure Goat Anti-Mouse IgG, Cy5-AffiniPure Donkey Anti-Guinea Pig IgG (1:200, Jackson ImmunoResearch). Confocal images were obtained with Leica SP5 confocal microscopy and analyzed by Leica software and Image J.

#### Quantification of fluorescence and statistical analysis

For lacZ/β-gal intensity analysis, including *EcRE-lacZ, Puc-lacZ* and *βFtz-f1-lacZ*, fluorescence intensity of border cell was measured by Image J and normalized to the nurse cells’ staining background to obtain the relative intensity. Statistical analysis was performed with GraphPad Prism 6 using unpaired two-tailed Student’s t-test, significance of p<0.05 was used as the criterion for statistical significance and indicated with *, p<0.01 was indicated with **, p<0.001 with three stars (***) and p<0.0001 with ****, not significant was indicated with “ns”.

#### 3-D imaging of border cell cluster

We used 2 coverslips (0.13-0.17mm thick) as bridges to mount egg chambers so that there is ample space in the z-axis to avoid compression of border cell clusters. Individual confocal sections were captured every 0.4 μm for each z-series of border cell cluster. The z-series was then processed by Imaris software to view 3-D distributions of aPKC, Lifeact-GFP, Par6-GFP and PIP2-GFP (Figure 3, Movies 1-4).

#### Chitin staining

Ovaries were fixed as described for immunostaining but without blocking. FB28 (Sigma) was used as a chitin dye as previously reported. We used FB28 (50mg/ml) with dilution of 1:400, stained ovaries in PBST for 30 min, washing 3×10 min with PBST.

